# Single-cell profiling of blood and cerebrospinal fluid in tuberculous meningitis

**DOI:** 10.1101/2025.03.04.641394

**Authors:** Trinh Thi Bich Tram, Lucy C. Garner, Le Nguyen Hong Thai, Le Thanh Hoang Nhat, Do Dang Anh Thu, Ho Dang Trung Nghia, Le Hong Van, Guy E Thwaites, Vu Thi Ngoc Ha, Paul Klenerman, Nguyen Thuy Thuong Thuong

## Abstract

Tuberculous meningitis (TBM) is the most severe form of tuberculosis, with a fatality rate of 20-50% in treated individuals. Although corticosteroid therapy can increase survival in HIV-negative people with TBM, better antimicrobial and host-directed therapies are required to improve outcome. There is, therefore, a need to better understand local immunopathologic pathways. Despite its power in identifying disease-specific cellular profiles, single-cell RNA-sequencing (scRNA-seq) has been underutilized in cerebral samples in brain infection. We employed scRNA-seq to analyze fresh pretreatment cerebrospinal fluid (CSF) from four TBM patients, along with paired peripheral blood mononuclear cells (PBMCs). While 29 cell subtypes were present in both tissues, their relative abundance varied significantly. In particular, CSF was enriched with highly inflammatory microglia-like macrophages, *GZMK*-expressing CD8^+^ T cells, and CD56^bright^ NK cells. The latter two subsets exhibited features associated with dysfunctional cytotoxicity. Across multiple cell types, inflammatory signaling pathways were increased and oxidative phosphorylation was decreased in CSF compared to PBMCs. This study highlights the value of scRNA-seq for exploring CSF immunopathogenesis in TBM patients and offers a resource for future studies investigating the pathophysiology of TBM and other brain infections, including potentially targetable cell populations linked with immune-mediated pathology.

## Introduction

Failures in the host response following inhalation of *Mycobacterium tuberculosis* (*Mtb*) can result in bacterial replication and hematogenous dissemination of bacteria to organs beyond the lung. Tuberculous meningitis (TBM) is one such example, accounting for approximately 5-10% of extra-pulmonary tuberculosis cases and 1% of active TB cases (1,2). TBM is the most severe and fatal form of TB, with a mortality rate of 20-50% in treated individuals and up to 50% long-term neurological disability among survivors (3,4).

Given the lack of direct access to brain tissue in people with TBM, cerebrospinal fluid (CSF) has been used as a proxy to study cellular and immunological events in the brain. CSF can be taken safely and repeatedly by lumbar puncture of the subarachnoid space. Immune responses in TBM were thought to be compartmentalized within the central nervous system, however, recent transcriptomic studies revealed significant levels of systemic inflammation (5–7). Whole blood RNA-sequencing (RNA-seq) of TBM patients with HIV coinfection showed increased neutrophil-associated transcripts and inflammasome signaling in those with immune reconstitution inflammatory syndrome compared with those without (5). T and B cell activation pathways were decreased in the blood of TBM patients with fatal outcomes, but their activities in CSF remain unknown (6). RNA-seq of lumbar CSF from children with TBM showed enhanced protein translation and cytokine signaling compared with other central nervous system (CNS) infections, while blood of TBM children exhibited upregulated inflammasome signaling compared with healthy donors (7). Given the localized CNS response in TBM, alongside known systemic effects, linking host immune responses in PBMCs and CSF is essential for understanding TBM pathogenesis.

While bulk RNA-seq provides an average gene expression profile of a mixed population of cells, single-cell RNA-seq (scRNA-seq) measures gene expression at the level of individual cells, enabling hypothesis-free identification and characterization of cell type-specific immune responses in human disease, including TB (8). In a previous scRNA-seq study, a cytotoxic NK cell subset (*CD7*^+^*GZMB*^+^) was depleted in PBMCs from latent TB compared with healthy individuals, and further depleted in active TB, suggesting a novel biomarker for distinguishing these manifestations (9). scRNA-seq of PBMCs from lung TB patients revealed expansion of exhausted Th1 cells, CD8^+^ T cells, and NK cells, with the latter two cell subsets exhibiting a highly cytotoxic phenotype in severe cases (10). Therefore, scRNA-seq offers a promising approach to characterize the CSF immunopathogenesis in TBM patients and could reveal novel cell or gene targets for TBM diagnostics and treatment.

Compared with blood, studying the CSF has several challenges, limiting its broad implementation in CNS infection research. Sampling is invasive, and samples are prone to blood contamination and typically contain low numbers of cells (7,11). Consequently, scRNA-seq on CSF in inflammatory diseases is typically performed with fresh samples (12–16). A previous study on two-month-frozen CSF samples showed the potential of using cryopreserved CSF in scRNA-seq, but further validation is required due to significant variation in cell viability and the number of cells recovered (17). Additionally, while comprehensive reference datasets of the blood transcriptional landscape in health and numerous diseases are publicly available, a reference dataset for CSF in adult TB is lacking. Herein we performed scRNA-seq on fresh paired PBMCs and CSF from four adult TBM patients. We identified 15 major cell types and 29 cell subtypes with distinct tissue distribution patterns. Furthermore, we revealed molecular changes in immunological and metabolic processes among PBMCs and CSF, as well as between TBM survivors and a non-survivor. Our findings collectively highlight the utility of scRNA-seq for providing deeper insights into TBM pathogenesis.

## Materials and Methods

### Participants

Of the four patients suspected of TBM, three were participants of the LAST ACT trial (NCT03100786), which evaluated the impact of adjunctive dexamethasone on outcomes for HIV-negative adults with TBM. This trial was conducted in the Hospital for Tropical Diseases (HTD) and Pham Ngoc Thach Hospital (PNT) in Ho Chi Minh City, Vietnam from February 2018 to March 2024 (18). The remaining patient was a participant of an ongoing cross-sectional study, started in May 2023 in the HTD, aimed at improving the diagnosis of CNS infectious diseases. Adults (≥ 18 years) who underwent lumbar puncture as part of routine care for suspected CNS infection were recruited. Patients were suspected of TBM if they had ≥ 5 days of meningitis symptoms and abnormal CSF parameters (including color, opening pressure, white blood cell count, protein, lactate, and glucose). One healthy donor was enrolled in an ongoing epidemiology study of human resistance to *Mtb* infection, conducted in the HTD in July 2022. Written informed consent was obtained from all participants or their relatives if they were incapacitated. Study protocols were approved by the HTD and PNT hospital in Vietnam, and the Oxford Tropical Research Ethics Committee, UK.

### Sample collection

As part of routine hospital procedure, 5-10 ml of CSF and 1 ml heparinized blood were collected. 2-3 ml of CSF were sent to the hospital laboratory for routine tests for viral, bacterial, or fungal infection. The remaining CSF was concentrated by centrifugation at 300 × g for 10 min and re-suspended in 700 µl of CSF supernatant. This CSF deposit was used for TB diagnostic tests, including 100 μl for Ziehl-Neelsen smear, 200 μl for Xpert MTB/RIF, and 200 μl was stored at −80°C for future use. The remaining 200 μl of CSF was used for scRNA-seq if processed within four hours of collection, with ≥ 75 cells/µl deposit, and no artificial blood contamination (evidenced by > 200 RBCs/μl CSF total) (15). After re-centrifugation (300 × g, 5 min), CSF pellets were resuspended in 50 μl PBS containing 0.04% bovine serum albumin (BSA, Sigma) and counted. Blood was processed alongside CSF, with PBMCs isolated using density gradient centrifugation with Histopaque 1077 (Sigma) following the manufacturer’s instructions. Both PBMCs and CSF were used fresh for downstream applications, including scRNA-seq and flow cytometry.

### Flow cytometry

PBMCs and CSF cells were stained with Zombie Aqua dye (BioLegend), then incubated with a cocktail of antibodies for 15 min at room temperature: CD3-PerCP (SK7), CD4-FITC (RPA-T4), CD8-APC/Cy7 (SK1), CD14-BV605 (63D3), CD16-PE/Cy7 (3G8), CD19-Alexa Fluor 700 (SJ25C1), CD56-BV421 (HCD56), CD123-PE (6H6), and CD11c-APC (SHCL-3). CD3-PerCP, CD4-FITC, and CD11c-APC were from BD Biosciences, while all other antibodies and dyes were from BioLegend. Samples were run on a BD FACSLyric flow cytometer using FACSuite acquisition software. Data was analyzed using FlowJo software (v10.9.0, BD) using the gating strategy depicted in **Figure S1**.

### Generation and sequencing of 10x Genomics scRNA-seq libraries

Single-cell suspensions were loaded onto a 10x Genomics Chromium Controller. One patient sample was loaded per channel with a target capture rate of 4,000 cells, while the healthy control PBMCs had a target capture rate of 1,000 cells. Libraries were generated using the Chromium Next GEM Single Cell 3’ Reagent Kits v3.1 according to the manufacturer’s instructions. The library from the healthy control PBMCs was sequenced internally on an Illumina MiSeq to a mean depth of 20,922 reads per cell, with a sequencing saturation of 66.1%. Patient libraries were sequenced on an Illumina NovaSeq 6000 (BGI, Hong Kong) to a mean depth of 33,072-163,448 reads per cell, with a sequencing saturation of 54.0-91.6%.

### scRNA-seq analysis

#### Pre-processing and quality control

Sequencing data was processed using Cell Ranger v7.1.0 (10x Genomics). Namely, reads were aligned to a GRCh38 reference genome (v3.0.0) and gene expression matrices generated using *cellranger count*. SoupX (v1.6.2) was used to remove ambient RNA contamination (19). Doublets were removed using Scrublet (v0.0.4) with default parameters (20). The data from all samples were combined using the merge function in Seurat v4.3.1 (21). Low quality cells with < 500 genes detected or > 10% of reads aligned to the mitochondrial genome were removed. Genes expressed in fewer than five cells were removed.

### Data normalization, dimensionality reduction, and clustering

After quality control, data was analyzed using Seurat v4.3.1 (21). Raw count data was normalized using sctransform (22) with percent mitochondrial reads and the number of unique molecular identifier (UMI) counts regressed out. Variance stabilizing transformation was performed to identify the 3,000 most variable genes, which were used for principal component analysis (PCA). Batch effects from different donors were removed using Harmony (23). A shared nearest neighbor graph was constructed using the top 30 Harmony components, and clustering was performed with the Louvain algorithm (resolution 1.9). Data was visualized using UMAP of the top 30 Harmony PCs.

For some analyses, we combined the data from TBM patients with that from one healthy donor. Raw counts from the healthy donor were subjected to the same quality control metrics as the TBM dataset before merging. The combined dataset was processed using the same procedures as for the TBM dataset.

### Cell annotations

Clusters were annotated at two levels: broad annotation of major cell types and deeper annotation of cell subtypes. For the first level, data was mapped to reference datasets of PBMCs and immune cells (Monaco and Blueprint/ENCODE) using Azimuth (24) and SingleR (25) algorithms, respectively. To improve annotation accuracy, we manually inspected the expression of known cell type marker genes. Additionally, we identified and reviewed the genes differentially expressed between clusters (*FindAllMarkers* function with MAST (26)). The MAST model included patient ID and cellular detection rate (CDR) as co-variates. Genes with a fold change ≥ 1.25 and a false discovery rate (FDR) < 0.05 were considered significantly enriched in a cluster.

In the combined dataset of cells from TBM patients and a healthy donor, cell types for the healthy donor were inferred by mapping to a reference of PBMCs from the four TBM patients using SingleR.

### Differential abundance analysis

To formally compare the cell type proportions among PBMCs and CSF, differential abundance testing was performed using miloR (v1.9.1) (27) with parameters k = 30, d = 30, and prop = 0.1. The design was ∼ tissue. The *plotDAbeeswarm* function was used with mixed populations removed. Neighborhoods with a spatial FDR < 0.1 were considered significantly differentially abundant.

### Differential expression analysis

Differentially expressed genes (DEGs) between PBMCs and CSF for each cell type, as well as DEGs between cell subtypes, were identified using the *FindMarkers* function from Seurat with MAST (26), including patient ID and CDR as co-variates. Cell types with < 10 cells were excluded from the analysis. Only genes detected in > 1% of cells in each cell type in a given tissue were included. Ribosomal and mitochondrial genes were excluded from the analysis. Significant genes were defined as those with a fold change ≥ 1.25 and an FDR < 0.05.

### Gene set enrichment analysis and over-representation analysis

Gene set enrichment analysis (GSEA) was conducted using clusterProfiler (v4.6.2) (28) with the fast gene set enrichment analysis (FGSEA) method (29). For each cell type, all genes were ranked in descending order by their log2 fold change values. “Hallmark” gene sets were obtained from the Molecular Signatures Database (MSigDB) (http://www.gsea-msigdb.org/gsea/index.jsp). Pathways with an FDR < 0.05 were considered significant. Over-representation analysis (ORA) for Gene Ontology terms was conducted using clusterProfiler (v4.6.2) with DEGs as input.

### Single-sample GSEA

To compare pathway activity between PBMCs and CSF of TBM patients or between TBM patients and a healthy control, pathway enrichment scores were calculated for single cells using the single-sample GSEA method implemented in GSVA (30). The Wilcoxon rank-sum test was used for comparison between groups. Gene sets associated with immune responses were compiled from the Gene Ontology database and the apoptosis pathway was from MSigDB, while metabolic pathways were from the Kyoto Encyclopedia of Genes and Genomes (KEGG) database.

### Data Availability Statement

The sequencing data generated in this paper have been deposited in NCBI’s Gene Expression Omnibus (GEO).

## Results

### Patient characteristics

To analyze the cell composition and transcriptional landscape of PBMCs and CSF in TBM, we performed scRNA-seq on fresh paired pretreatment samples from four patients (**Figure 1A**). Baseline characteristics of the patients are summarized in **Table 1**. All four patients were clinically diagnosed with TBM, with microbiological confirmation in two cases. White blood cells ranged from 7,000 to 11,000 cells per ml, with neutrophils comprising the largest proportion (∼80%), followed by lymphocytes. All four patients exhibited a high white cell count in CSF (∼300,000 cells/ml), with lymphocytes being the predominant cell type (> 80%). Three patients survived, whereas one with a low CSF white cell count died within a week of hospitalization.

**Figure 1.**
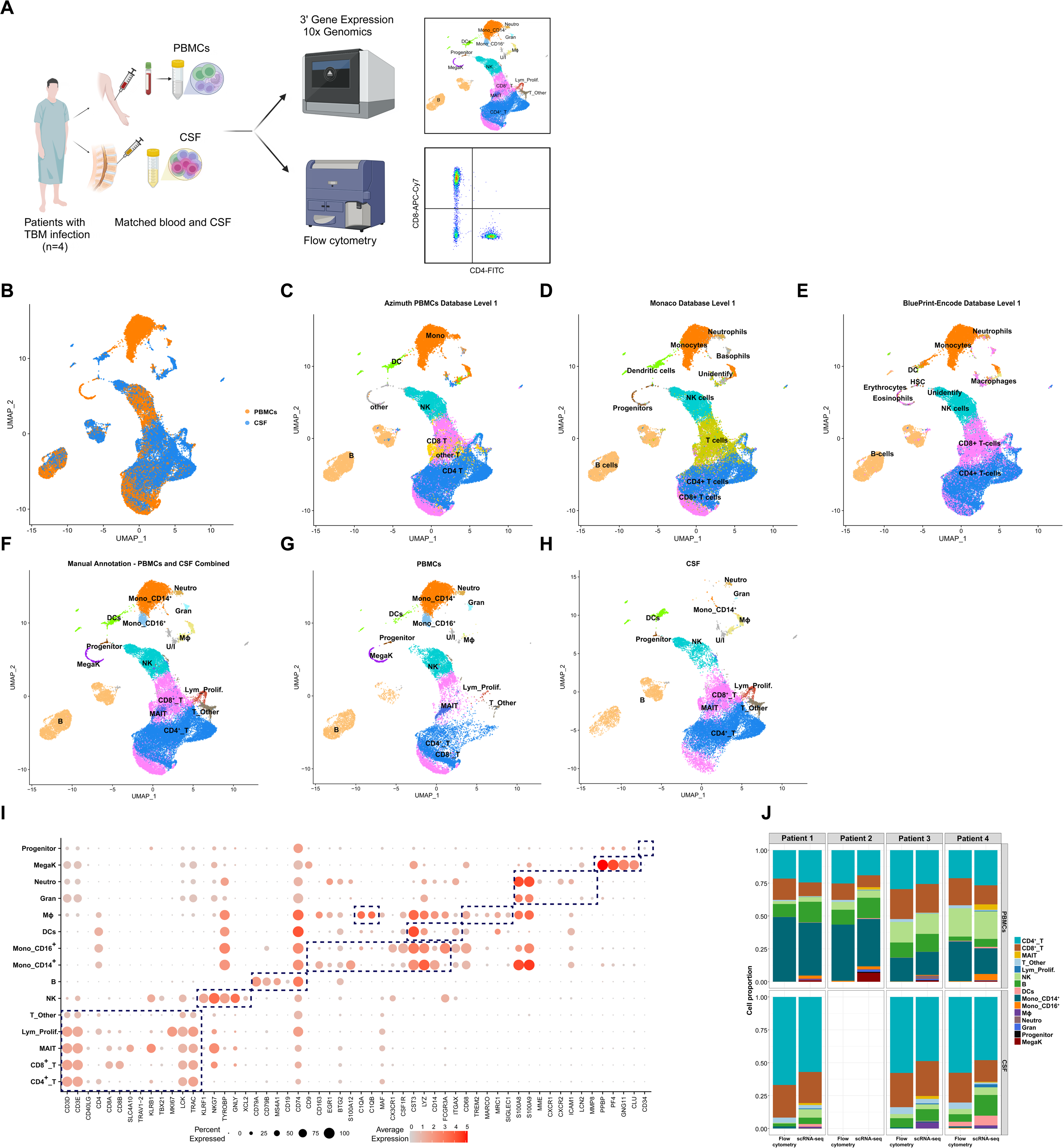
Main cell types (Level 1) in combined PBMCs and CSF, and comparison with flow cytometry results. (**A**) Experimental workflow. Figure generated using BioRender. (**B**) UMAP plot of all cells colored by tissue of origin. (**C-E**) UMAP showing Level 1 annotation of cell types for combined PBMCs and CSF based on reference datasets: (**C**) Azimuth for PBMCs, (**D**) Monaco for immune cells, and (**E**) Blueprint/ENCODE for stromal and immune cells. (**F-H**) UMAP showing Level 1 annotation of cell types based on expert knowledge for: (**F**) combined PBMCs and CSF or separated into (**G**) PBMCs and (**H**) CSF. (**I**) Dot plot depicting selected marker genes of major cell types (indicated by dashed boxes). Dot color indicates the mean normalized expression and dot size indicates the fraction of cells expressing the gene. (**J**) Comparison of cell proportions identified by flow cytometry and scRNA-seq. B: B cells, CD4^+^_T: CD4^+^ T cells, CD8^+^_T: CD8^+^ T cells, DCs: dendritic cells, Gran: mixed granulocytes, Lym_Prolif.: proliferative lymphocytes, MAIT: MAIT cells, MegaK: megakaryocytes, Mono_CD14^+^: CD14^+^ monocytes, Mono_CD16^+^: CD16^+^ monocytes, MΦ: macrophages, Neutro: neutrophils, NK: NK cells, T_Other: other T cells, U/I: unidentified.

**Table 1.** Baseline characteristics of four TBM patients.

| Characteristics | Patient 1 | Patient 2 | Patient 3 | Patient 4 |
| --- | --- | --- | --- | --- |
| Age (years) | 40 | 60 | 40 | 51 |
| Gender | Male | Male | Male | Male |
| Ziehl-Neelsen stain | Positive | Negative | Negative | Negative |
| Xpert Ultra | Trace | Low | Negative | Negative |
| MGIT culture | Negative | Positive | Negative | Not done |
| HIV infection | Negative | Negative | Negative | Negative |
| Cryptococcal antigen LFA | Negative | Negative | Negative | Negative |
| Diagnostic category* | Definite TBM | Definite TBM | Probable TBM | Possible TBM |
| MRC grade** | Grade 2 | Grade 2 | Grade 1 | Grade 1 |
| Blood leukocytes ($10^6$ cells/ml) | 11.4 | 7.5 | 11.3 | 7.8 |
| Blood monocyte (%) | 4.0 | 5.4 | 4.9 | 8.2 |
| Blood lymphocyte (%) | 12.0 | 10.0 | 13.0 | 27.4 |
| Blood neutrophil (%) | 79.9 | 84.4 | 81.7 | 56.4 |
| Blood eosinophil (%) | 2.0 | 0.1 | 0.3 | 5.2 |
| CSF leukocytes ($10^3$ cells/ml) | 382.0 | 140.0 | 330.0 | 357.0 |
| CSF neutrophil (%) | 13.0 | 0.0 | 0.0 | 8.0 |
| CSF Lymphocyte (%) | 87.0 | 100 | 100 | 85.0 |
| CSF Eosinophil (%) | 0.0 | 0.0 | 0.0 | 0.0 |
| Treatment response at hospital discharge | Survived | Died within 1 week of hospitalization | Survived | Survived |

\*Diagnostic categories were assigned according to the consensus case definition (31).
\*\*MRC grade denotes modified British Medical Research Council criteria
LFA: Lateral flow assay; TBM: Tuberculosis meningitis

### Broad annotation (Level 1) of immune cell types in blood and CSF

A total of 19,921 and 21,292 single-cell transcriptomes were obtained from matched PBMCs and CSF, respectively. The number of genes detected per cell was comparable between PBMCs and CSF (2,723 ± 226 and 2,773 ± 550, respectively) (**Table S1**). Data was of a high quality, with low percentages of doublets (< 5%), ambient RNA (0.02%), and cells with > 10% mitochondrial reads (< 2%) (**Figure S2**). After removal of low-quality cells, 17,264 PBMCs and 20,100 CSF cells were combined for analysis (**Figure 1B**).

PBMCs and CSF cells partially overlapped on the Uniform Manifold Approximation and Projection (UMAP) (**Figure 1B**), suggesting a mixture of shared and compartment-specific cell types. As there is no reference CSF database for cell annotation, we reference mapped combined PBMCs and CSF to the Azimuth PBMCs reference using Azimuth (**Figure 1C**), and to Monaco and Blueprint/ENCODE references using SingleR (**Figure 1D, E**). The number of detected cell types varied with each reference: Azimuth annotated seven major cell types while the others annotated eleven. Common clusters included B cells, CD4^+^ T cells, CD8^+^ T cells, monocytes, and natural killer (NK) cells. However, inconsistencies in annotations were observed. For instance, Monaco identified progenitors, whereas Blueprint/ENCODE labeled the same cells as eosinophils and erythrocytes, and Azimuth did not recognize them. Additionally, due to the lack of neutrophils in PBMCs, Azimuth was unable to annotate neutrophils. To reconcile annotation differences, we further manually annotated cell types using expert knowledge (**Table S2**). Louvain clustering at resolution 1.9 produced 41 clusters (**Figure S3**) which were subsequently classified into 15 major cell types based on canonical marker gene expression (**Figure 1F-I**). Identified cell types included B cells (*CD79A*/*B*, *MS4A1*, *CD74*, *CD19*), CD4^+^ T cells (*CD3D*/*E*, *CD4*), CD8^+^ T cells (*CD3D*/*E*, *CD8A*/*B*), mucosal-associated invariant T (MAIT) cells (*SLC4A10*), proliferative lymphocytes (*MKI67*), “T_Other” cells (possibly dead cells due to low RNA content, **Figure S4**), NK cells (*KLRF1*, *GNLY*, *XLC2*), dendritic cells (DCs) (*CST3*, *LYZ*, *ITGAX*), two types of monocytes (CD14^+^ [*CD14*] and CD16^+^ [*FCGR3A*]), macrophages (*C1QA*, *C1QB*, *MARCO*, *SIGLEC1*, *TREM2*, *MRC1*), megakaryocytes (*PPBP*, *PF4*, *GNG11*, *CLU*), progenitors (*CD34*), and two granulocyte types (neutrophils: *S100A8*, *S100A9*, *ICAM1*, *CXCR2*, *MME*; mixed granulocytes: *MMP8*, *LCN2*). All cell types were present in both PBMCs and CSF, except for megakaryocytes and CD16^+^ monocytes, which were only found in blood (**Figure 1G, H**).

We validated the cell composition identified using scRNA-seq with flow cytometry (**Figure 1J**). Flow cytometry detected CD4^+^ T cells, CD8^+^ T cells, and other CD3^+^ T cells. As anticipated, scRNA-seq provided a higher resolution of cell subtypes compared with flow cytometry, but there was high concordance in the frequencies of shared cell types. Overall, microfluidics-based scRNA-seq successfully characterized the cell composition of PBMCs and CSF from adult TBM patients.

### High-resolution annotation (Level 2) of immune cell subtypes in blood and CSF

At the broad level of annotation, multiple Louvain clusters were identified within B cells, NK cells, CD4^+^ T cells, CD8^+^ T cells, and DCs, suggesting the presence of cell subtypes (**Figure 1F, Figure S3**). We further annotated the subtypes based on marker gene expression (**Figure 2A, B**). This considerably increased cell annotation granularity to 29 subtypes (**Figure 2A**). For example, we identified four subsets of B cells: naïve (*TCL1A*, *IGHD*), memory (*JAM3*), plasmablast (*CD27*, *CD38*, *XBP1*, *MZB1*, *MKI67*), and plasma cells (similar to plasmablast, but negative for *MKI67*). CD4^+^ T cells divided into three subtypes: CD4^+^ naïve/T central memory (TCM) (*CCR7*, *SELL*), Treg (*FOXP3*, *IL2RA*), and CD4^+^ memory (CD4^+^ TM) (*GPR183*, *S100A4*). CD8^+^ T cells comprised four clusters: CD8^+^ naïve/TCM (*CCR7*, *SELL*) and three effector-memory populations, namely GNLY^+^CD8^+^ TEM (high *GNLY*), GZMK^+^CD8^+^ TEM (high *GZMK*, low *GNLY*), and activated GZMK^+^CD8^+^ TEM (high inhibitory receptors: *TIGIT*, *LAG3*, *CTLA4*, *HAVCR2*). DCs included three subtypes: conventional DC1 (cDC1; *ITGAX*, *WDFY4*, *XCR1*, *BATF3*), conventional DC2 (cDC2; *ITGAX*, *FCER1A*, *CLEC10A*), and plasmacytoid DCs (pDCs; *LILRA4*, *ITM2C*). Three subtypes of NK cells were identified: immunomodulatory CD56^bright^ NK (high *NCAM1*, *CD160*, *XCL1*), cytotoxic CD56^dim^ NK (moderate *NCAM1*, *CD160*), and adaptive-like NK (32,33) (moderate *NCAM1*, *B3GAT1*, *LILRB1*, very low *KLRC1*). Macrophages were divided into two clusters: microglia-like macrophages (high *C1QA*/*B*/*C*, *APOE*, *MRC1*, *TLR2*, *TLR4*) and macrophages (low *C1QA*/*B*/*C*). Except for monocytes and rare cell types like cDC2, most subtypes were present in PBMCs and CSF in all four patients (**Figure S5**). Overall, CSF comprised a large variety of cell subtypes, with comparable diversity to blood.

**Figure 2.**
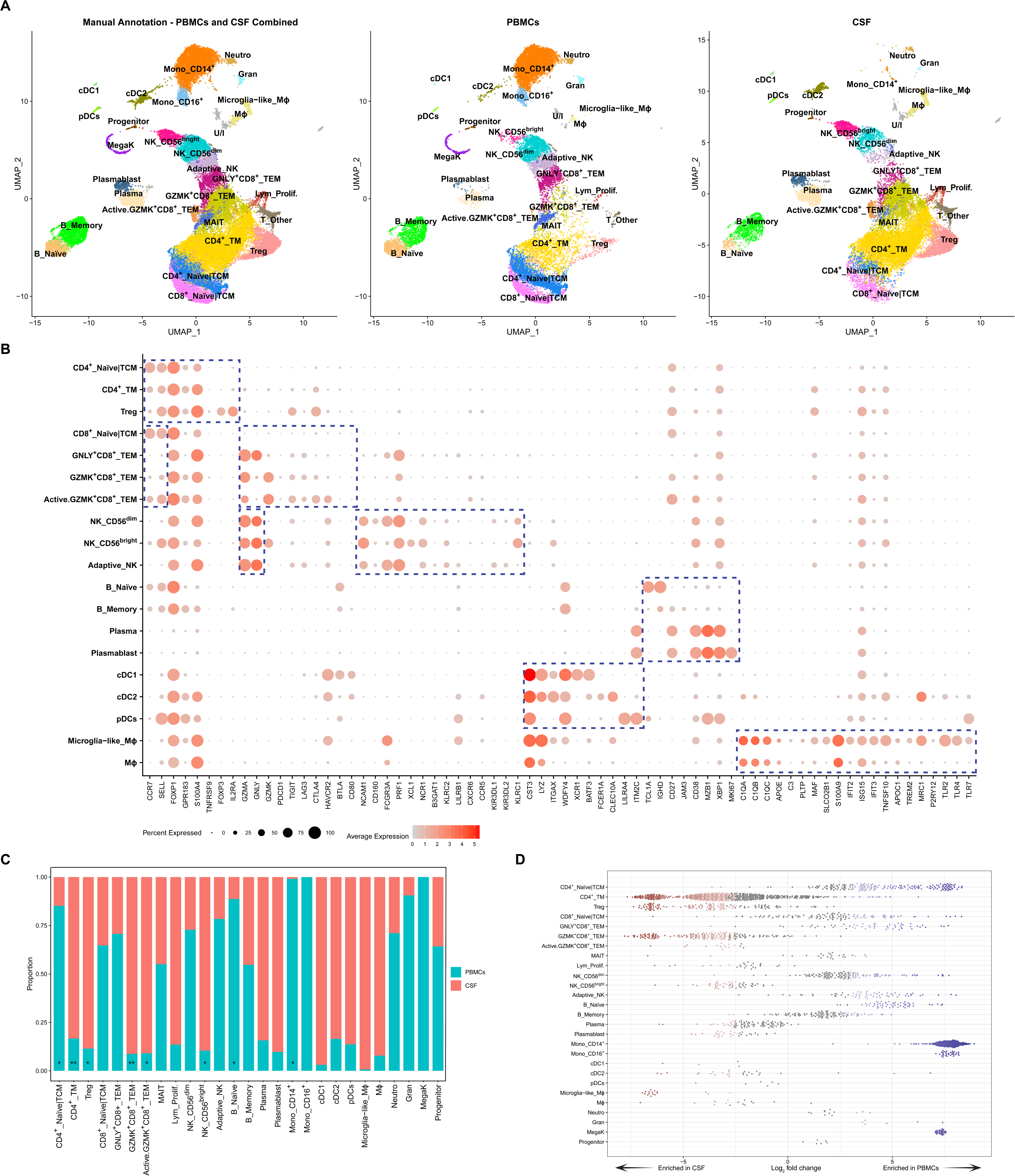
Cell subtypes (Level 2) in combined PBMCs and CSF. **(A)** UMAP plot of high-resolution annotation (Level 2) of cell subtypes from the main cell types, based on expert knowledge for combined PBMCs and CSF cells (left), PBMCs (middle), or CSF cells (right). **(B)** Dot plot depicting selected signature genes of cell subtypes (indicated by dashed boxes). Dot color indicates the mean normalized expression and dot size indicates the fraction of cells expressing the gene. (**C**) The proportions of cell subtypes in PBMCs and CSF cells from all four TBM patients. Comparison of cell subtype proportions between PBMCs and CSF across four patients was performed using paired t-tests. *p < 0.05; **p < 0.01. (**D**) Relative abundance of neighborhoods identified using the Milo k-nearest neighbor weighted-network algorithm. Neighborhoods enriched in CSF are in red, those enriched in PBMCs are in blue, and those with no significant enrichment are in grey. Adaptive_NK: adaptive-like NK cells, cDC1: conventional DC1, cDC2: conventional DC2, Gran: mixed granulocytes, Lym_Prolif.: proliferative lymphocytes, MegaK: megakaryocytes, MΦ: macrophages, Neutro: neutrophils, pDCs: plasmacytoid DCs, TCM: T central memory cells, TEM: T effector memory cells, TM: T memory cells, U/I: unidentified.

### Relative abundance of cell types in blood and CSF

We next analyzed differences in cell composition between PBMCs and CSF, focusing on the 27 cell subtypes annotated at high resolution. The unidentified group and “T_Other” (**Figure S4**) were excluded from the analysis. As expected, megakaryocytes were absent in CSF. CD14^+^ monocytes and CD16^+^ monocytes were almost exclusive to PBMCs, while microglia-like macrophages were mainly in CSF (**Figure 2C**). The following cell clusters predominantly comprised cells from CSF in all patients: Treg (p = 0.01), CD4^+^ TM (p = 0.008), GZMK^+^CD8^+^ TEM (p = 0.009), activated GZMK^+^CD8^+^ TEM (p = 0.003), and NK CD56^bright^ (p = 0.03). In contrast, CD4^+^ naïve/TCM (p = 0.01) and naïve B (p = 0.04) were over-represented in PBMCs (**Figure 2C, Figure S6**).

Limitations in defining the appropriate cell clustering resolution for annotation can hinder the identification of condition-specific cell subtypes or cell states (27). Therefore, we performed formal differential abundance analysis to compare cell proportions in PBMCs and CSF within local neighborhoods using the Milo k-nearest neighbor (k-NN) approach. Among 2,896 neighborhoods spanning the k-NN graph generated from PBMCs and CSF cells, Milo identified 1,845 neighborhoods with differential abundance (spatial FDR < 10%) (**Figure S7**). Consistent with the annotated cell type-based analysis, neighborhoods of microglia-like macrophages, CD4^+^ TM, and Treg cells were enriched in CSF (**Figure 2D**). Conversely, neighborhoods of monocytes (CD14^+^ and CD16^+^), naïve/TCM cells (CD4^+^ and CD8^+^), and naïve B cells were enriched in PBMCs. Notably, GZMK^+^CD8^+^ TEM, activated GZMK^+^CD8^+^ TEM, and CD56^bright^ NK cells were almost exclusive to CSF. In contrast, GNLY^+^CD8^+^ TEM, adaptive-like NK, and CD56^dim^ NK cells were abundant in blood and relatively depleted in CSF. Some neighborhoods of plasma cells, plasmablasts, and subsets of DCs were enriched in CSF. Collectively, these data demonstrate highly compartment-specific cell compositions.

We further investigated site-specific differences in effector-memory CD8^+^ T cells and NK cells. A total of 360 genes were differentially expressed between GZMK^+^CD8^+^ TEM and GNLY^+^CD8^+^ TEM cells (**Table S5**). Among these, the CSF-exclusive GZMK^+^CD8^+^ TEM population significantly upregulated granzyme K (*GZMK*), markers for naïve/central memory T cells (*SELL*, *TCF7*, *CCR7*), co-stimulatory receptors (*CD27*, *CD28*, *ICOS*), cytokines (*LTB*), cytokine and chemokine receptors (*IL7R*, *IL12RB2*, *CXCR3*/*4*/*6*), activation markers (*CD38*, *CD69*), and IFN-stimulated genes (*STAT1*, *ISG20*, *IRF1*). In contrast, genes encoding cytotoxic molecules (*GZMB*/*H*, *GNLY*, *PRF1*), cytotoxicity-related genes (*FGFBP2*, *NKG7*), and markers associated with effector functions and terminal differentiation (*CX3CR1*, *TBX21* [encoding T-bet], *ZEB2*) were upregulated in GNLY^+^CD8^+^ TEM cells (**Figure 3A, Figure S8**). CD56^bright^ NK cells differentially expressed 605 genes compared with non-CD56^bright^ NK cells (including both CD56^dim^ and adaptive-like NK cells) (**Table S6**), 179 of which overlapped with the differentially expressed genes (DEGs) between GZMK^+^CD8^+^ TEM and GNLY^+^CD8^+^ TEM (**Figure 3B, C**). Similar to GZMK^+^CD8^+^ TEM cells, CD56^bright^ NK cells showed upregulation of *GZMK,* naïve/central memory T cell markers (*SELL*, *TCF7*, *CCR7*), cytokines (*LTB*), and cytokine and chemokine receptors (*IL7R*, *IL12RB2*, *IL18R1*, *CXCR3*) (**Figure 3B, Figure S8**). In contrast, non-CD56^bright^ NK cells, enriched in PBMCs, expressed cytotoxicity-related genes (*GZMB*/*H*/*M*, *PRF1*, *FGFBP2*, *NKG7*), and genes associated with effector differentiation (*CX3CR1, TBX21*, *ZEB2*) (**Figure S8**), analogous to the GNLY^+^CD8^+^ population. Over-representation analysis revealed an enrichment of Gene Ontology terms related to cell differentiation and cytokine-mediated signaling in GZMK^+^CD8^+^ TEM and CD56^bright^ NK cells, while cytotoxicity/cell killing pathways were upregulated in GNLY^+^CD8^+^ TEM and non-CD56^bright^ NK cells (**Figure 3D-G**).

**Figure 3.**
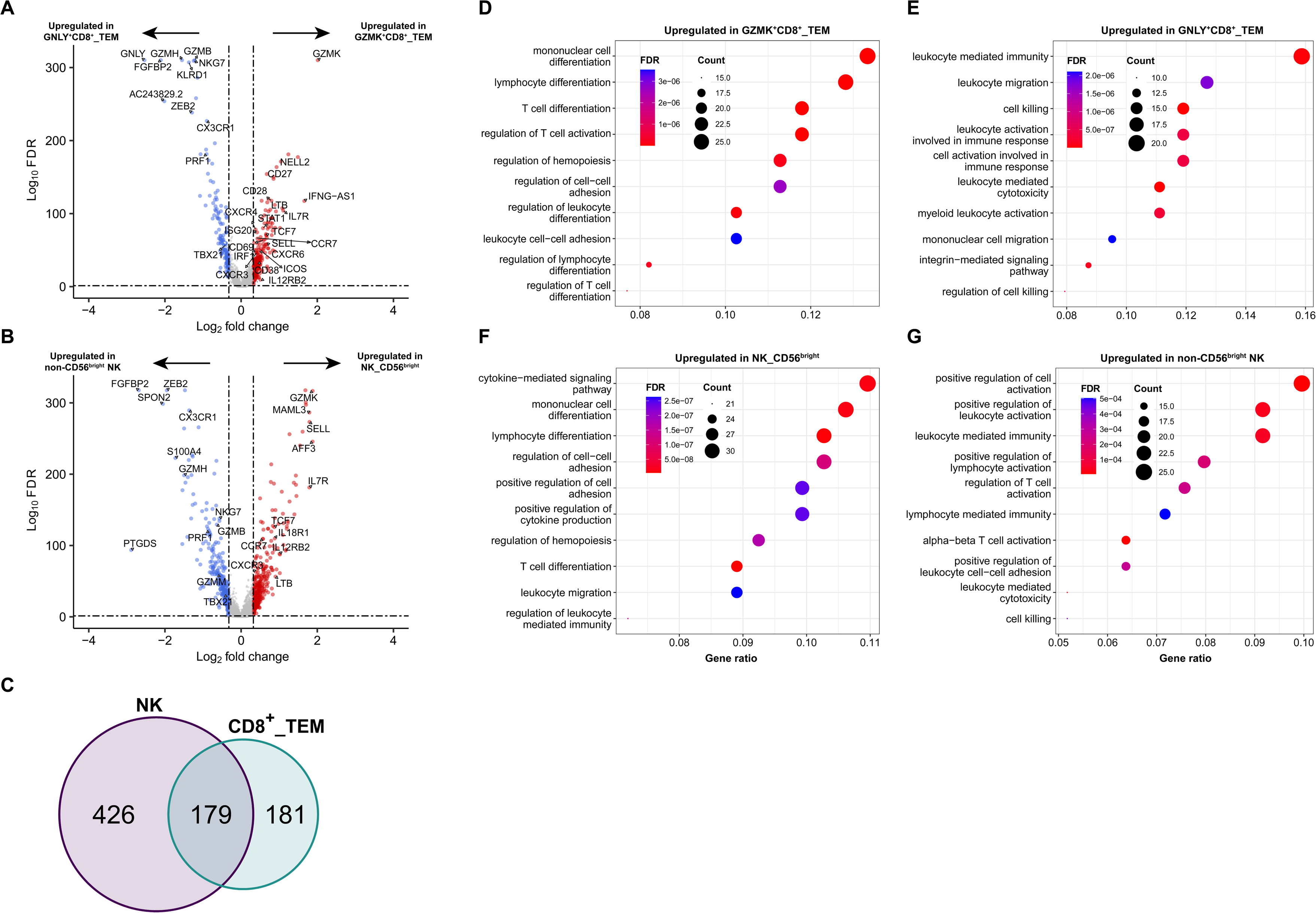
Differential expression of genes and pathways in effector memory CD8^+^ T cells and NK cells. (**A, B**) Volcano plot showing differentially expressed genes (DEGs) between (**A**) GZMK^+^CD8^+^ TEM and GNLY^+^CD8^+^ TEM cells, and between (**B**) CD56^bright^ NK and non-CD56^bright^ NK cells (including CD56^dim^ and adaptive-like NK cells). Top 10 and additional example DEGs are labeled. (**C**) Number of unique and common DEGs between the two effector memory CD8^+^ T cell populations and the two NK cell populations. (**D, E**) Over-representation analysis (ORA) for upregulated genes in (**D**) GZMK^+^CD8^+^ TEM and (**E**) GNLY^+^CD8^+^ TEM cells. (**F, G**) ORA for upregulated genes in (**F**) CD56^bright^ NK and (**G**) non-CD56^bright^ NK cells. Top 10 upregulated Gene Ontology Biological Process terms are shown. Gene ratio indicates the fraction of DEGs in the gene set.

### Enrichment of activation pathways and altered metabolism in CSF compared with PBMCs

We next determined compartment-specific transcriptional differences at the cell subtype level by identifying DEGs between PBMCs and CSF using MAST (26). Since CD16^+^ monocytes and microglia-like macrophages were found exclusively in PBMCs and CSF, respectively (**Figure 2C, D**), analysis could not be performed for these clusters. Moreover, due to the significantly lower number of CD14^+^ monocytes in CSF compared with PBMCs (40 vs. 4,230), we did not perform differential expression analysis for this cluster. Of the 29,000 genes detected, 1,834 (6.3%) genes were significantly differentially expressed (FDR < 0.05, FC ≥ 1.25) between PBMCs and CSF (**Figure 4A**). In general, more genes were upregulated in CSF compared with PBMCs, suggesting increased cell activity in CSF (**Figure 4A**). The rare cDC2 subset had the largest number of DEGs, with 500 genes upregulated in CSF and 400 genes in PBMCs. Integrated analysis revealed 408 universal DEGs (differentially expressed in ≥ five cell types) (**Table S3**) and 1,426 cell subtype-specific genes (**Table S4**). Universal DEGs upregulated in CSF were involved in lymphocyte migration (*CXCR6*) and activation (*CD69*), type I and II interferon responses (*STAT1*, *IRF1*, *SOCS1*, *GBP1*/*2*/*4*/*5* for both; *ISG15*, *ISG20* for type I interferon), neuroinflammation and neurodegeneration (*CD38*) (34), cytotoxicity (*APOL6* (35), *GZMB*), and T cell activation/exhaustion (*LAG3*) (**Table S3**). Universal DEGs upregulated in PBMCs were associated with cell proliferation and polarization (*RIPOR2*), migration (*TAGLN2*, *CRIP1*), cell activation and differentiation (*KLF2*, *CD52*), metabolite exchange (*UCP2*), and inflammation and tissue repair (*S100A4*/*10*) (**Table S3**). These findings reveal compartment-specific transcriptional landscapes in PBMCs and CSF, alongside differences in cell composition.

**Figure 4.**
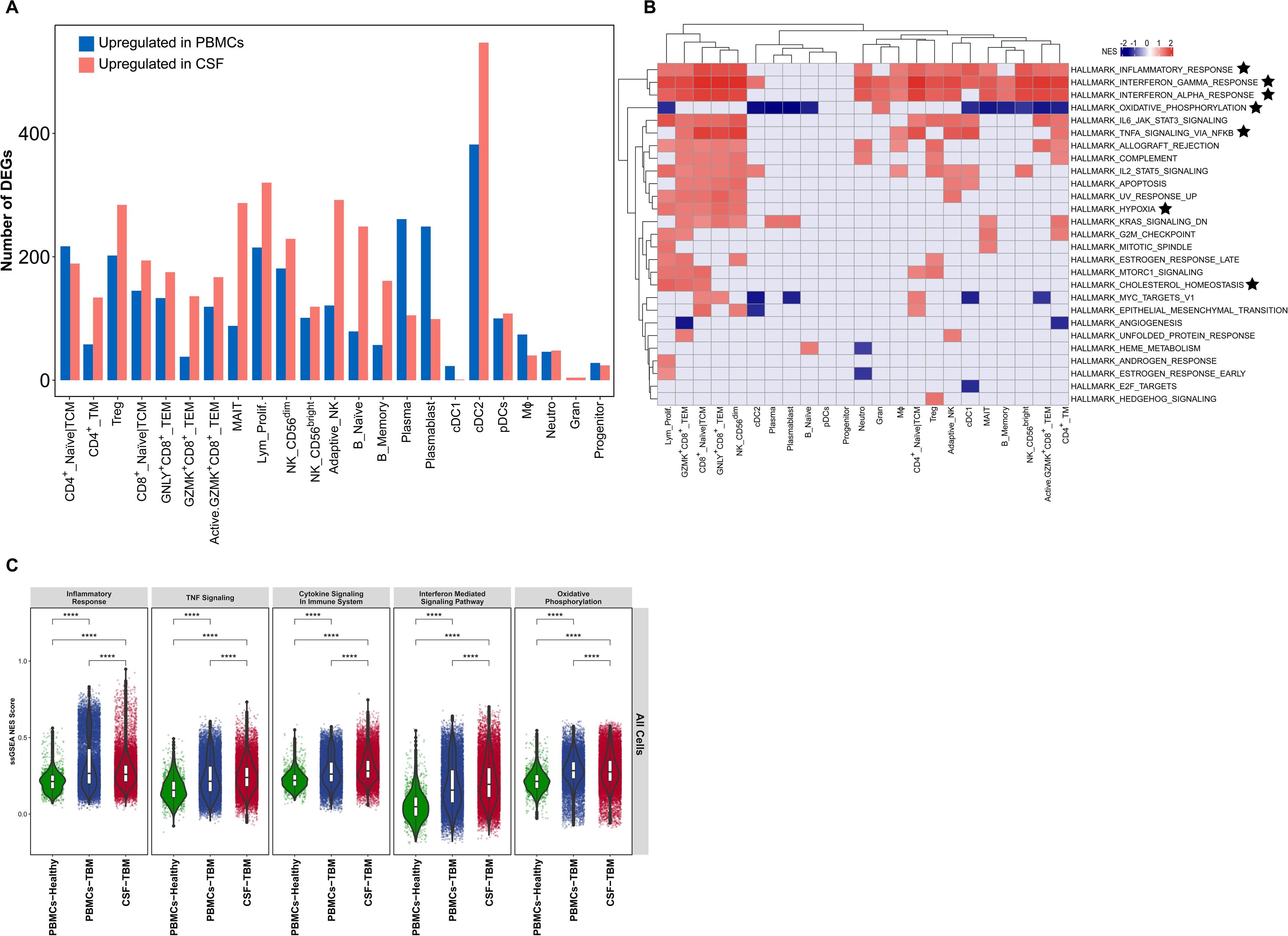
Differential expression of genes and pathways in individual cell subtypes from PBMCs and CSF of TBM patients. (**A**) Number of upregulated genes in individual cell subtypes in PBMCs and CSF. (**B**) Fast gene set enrichment analysis (FGSEA) for each cell subtype. Heatmap shows shared differentially active MSigDB Hallmark gene sets. Red indicates upregulation in CSF, blue indicates upregulation in PBMCs, and grey indicates no significant difference between PBMCs and CSF. Stars indicate gene sets discussed in the text. cDC1: conventional DC1, cDC2: conventional DC2, Gran: mixed granulocytes, Lym_Prolif.: proliferative lymphocytes, MegaK: megakaryocytes, MΦ: macrophages, Neutro: neutrophils, pDCs: plasmacytoid DCs, TCM: T central memory cells, TEM: T effector memory cells, TM: T memory cells. (**C**) Pathway normalized enrichment scores (NES) were calculated for individual cells using single-sample GSEA (ssGSEA). Each dot represents one cell. Boxes indicate the median, and 25^th^ and 75^th^ percentiles, with whiskers extending to the minimum and maximum data points. Statistical comparison between groups was performed using the Wilcoxon rank-sum test. ****p < 0.0001.

To further characterize the DEGs, we performed functional enrichment analysis for MSigDB Hallmark gene sets using fast gene set enrichment analysis (FGSEA) (29) (**Figure 4B**). Consistent with the universal DEGs, gene sets associated with inflammation and interferon alpha/gamma (type I/II) responses were enriched in multiple subsets of T and NK cells, and in macrophages, in the CSF. Similarly, the TNF signaling pathway was upregulated in the CSF in multiple subsets of T and NK cells, and in cDC1s. Intriguingly, hypoxia and cholesterol homeostasis pathways were enriched in several cell types in CSF, while oxidative phosphorylation was universally reduced. FGSEA results were confirmed with single-sample GSEA (ssGSEA), showing enrichment of total cytokine signaling, interferon signaling, and TNF signaling in CSF compared with PBMCs (**Figure 4C, Figure S9**). Compared to the healthy donor, TBM patients showed increased activity of cytokine response, inflammatory signaling, and oxidative phosphorylation pathways in both PBMCs and CSF (**Figure 4C**).

Among the four TBM patients, one succumbed to the disease. PBMCs and CSF from the non-survivor were enriched (heterogeneously across cell types) for a number of Hallmark gene sets associated with the immune response and metabolism. Namely, in PBMCs, activity of inflammatory, and interferon gamma and TNF signaling pathways, was increased in subsets of T cells, NK cells, and monocytes (**Figure 5A**). Enrichment of these pathways, along with the interferon alpha response, was also observed across multiple cell types in the CSF (**Figure 5B**). The oxidative phosphorylation pathway showed increased activity in most cell types in PBMCs and CSF from the non-survivor, indicating metabolic changes associated with TBM mortality.

**Figure 5.**
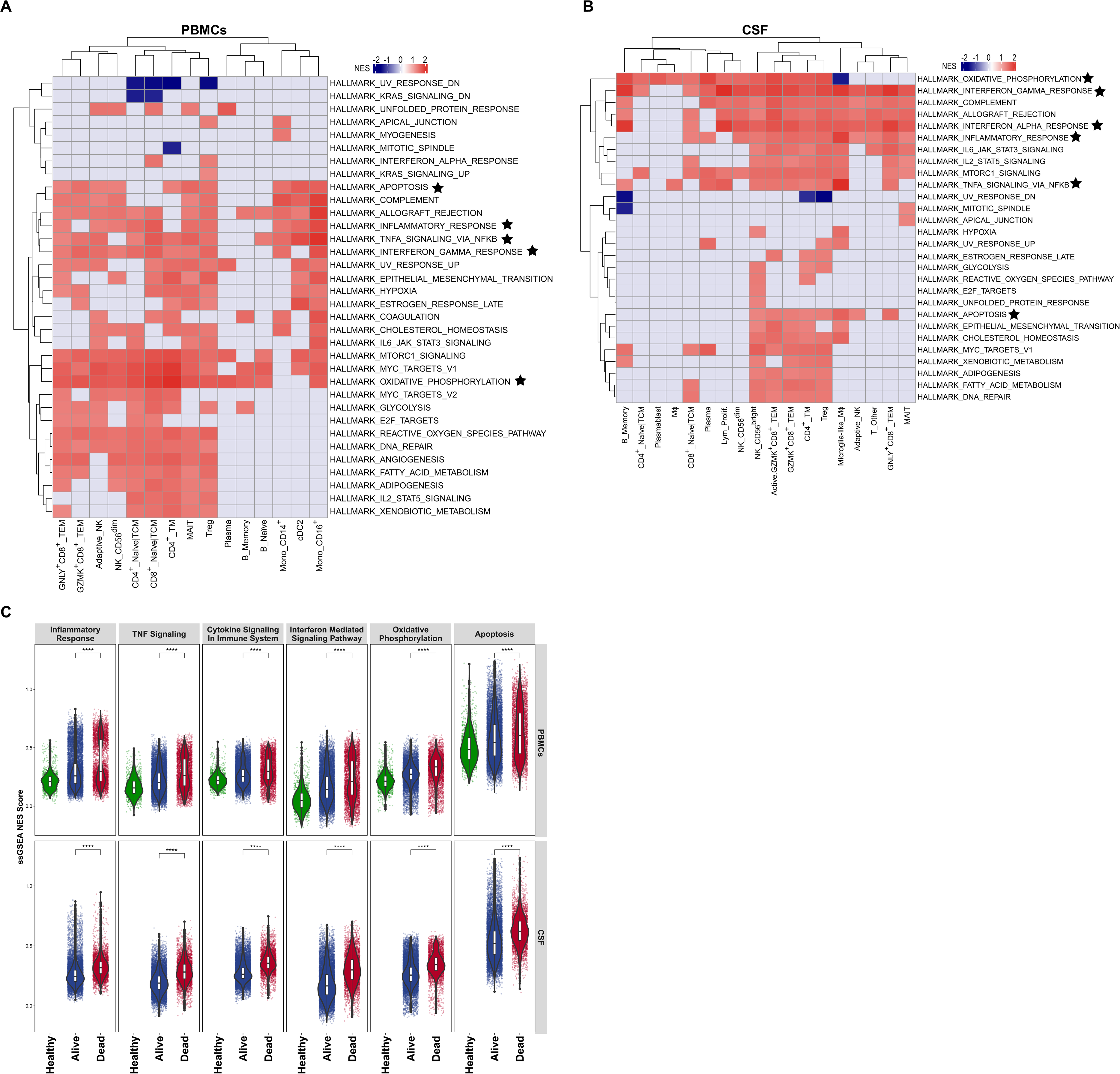
Fast gene set enrichment analysis (FGSEA) in individual cell subtypes from survivors and a non-survivor TBM patient. **(A, B)** FGSEA for each cell subtype in (**A**) PBMCs or (**B**) CSF between survivors and a non-survivor. Heatmap shows shared differentially active MSigDB Hallmark gene sets. Red indicates upregulation in the non-survivor, blue indicates upregulation in survivors, and grey indicates no significant difference between survivors and the non-survivor. Stars indicate gene sets discussed in the text. cDC2: conventional DC2, Lym_Prolif.: proliferative lymphocytes, MΦ: macrophages, TCM: T central memory cells, TEM: T effector memory cells, TM: T memory cells. (**C**) Pathway normalized enrichment scores (NES) were calculated for individual cells using single-sample GSEA (ssGSEA). Each dot represents one cell. Boxes indicate the median, and 25^th^ and 75^th^ percentiles, with whiskers extending to the minimum and maximum data points. Statistical comparison between groups was performed using the Wilcoxon rank-sum test. ****p < 0.0001.

The apoptosis gene set was enriched in several subsets of T cells, NK cells, and monocytes from the non-survivor, with highest activity in GNLY^+^CD8^+^ TEM, GZMK^+^CD8^+^ TEM, adaptive-like NK, CD4^+^ TM, MAIT, CD14^+^ monocytes and CD16^+^ monocytes in PBMCs, and in highly inflammatory CD56^bright^ NK, GZMK^+^CD8^+^ TEM, activated GZMK^+^CD8^+^ TEM, GNLY^+^CD8^+^ TEM and microglia-like macrophages in CSF (**Figure 5A, B**). Increased activity of these pathways in multiple cell types was consistent with their higher overall activity, as measured by ssGSEA (**Figure 5C**).

## Discussion

Over the past decade, significant insights have been gained into the cellular and biochemical environment of CSF in CNS infection, particularly in TBM, contributing to advances in diagnosis and treatment. However, the complex biological processes and disease mechanisms remain poorly defined, necessitating high-resolution tools to capture the full immune landscape in the CSF. For the first time, we employed scRNA-seq in paired PBMCs and CSF from adults with TBM. We demonstrated the feasibility of using scRNA-seq in CSF, characterized the cellular composition in CSF, and uncovered distinct yet related cellular and gene expression profiles in PBMCs and CSF in TBM.

We standardized protocols for studying CSF using scRNA-seq in multiple ways. Firstly, we paired CSF with matched PBMCs from each patient, using the PBMCs as a reference. Second, we implemented a two-step annotation process to confidently classify cell types. Third, we confirmed consistent major cell type proportions with flow cytometry. In total, we identified 15 major cell types and 29 cell subtypes in CSF, demonstrating a diversity comparable to that observed in PBMCs.

Single-cell transcriptional characterization of CSF from TBM patients showed differences with studies of other CNS diseases. In bacterial meningitis, neutrophils and monocytes were the main populations in the CSF (14). In contrast, CSF from COVID-19 patients was rich in T cells and CNS-associated macrophages (36). In multiple sclerosis, T cells made up the majority of cells in the CSF (15). In a previous study of children with TBM (37), CSF predominantly comprised three main populations: T cells (CD4^+^ and CD8^+^), B cells, and NK cells, consistent with our data. Thus, different cell types are mobilized to the CNS depending on the neuroinflammatory disorder.

In this study, we identified an abundance of *GZMK*-expressing CD8^+^ TEM and CD56^bright^ NK cell populations in the CSF compared with PBMCs. GZMK^+^CD8^+^ TEM are enriched in inflamed tissues, including CSF from children with TBM (37), pleural fluid from patients with tuberculous pleural effusion (38), and gut from patients with Crohn’s disease and ulcerative colitis (39). CD56^bright^ NK cells were the most abundant NK cell population in the CSF of TBM patients (40) and in the CNS in multiple CNS disorders (41). While blood-abundant GNLY^+^CD8^+^ TEM, CD56^dim^ NK, and adaptive-like NK cells exhibited a classic cytotoxic phenotype in the CSF of TBM patients, GZMK^+^CD8^+^ TEM and CD56^bright^ NK cells showed low cytotoxic potential and enriched activity of cytokine-mediated signaling pathways. Such differences in the abundance of *GNLY*- and *GZMK*-expressing populations in PBMCs and CSF suggested a transition in cell phenotype driven by the local environment. Granzyme K is a serine protease released by cytotoxic lymphocytes that may induce apoptosis and contributes to the pathogenesis of inflammatory skin diseases, viral infections, and sepsis (42–44). Granzyme K itself could act as a key inflammatory factor by activating pro-inflammatory pathways, for example production of IL-6 and CCL2 (39). Together, these findings suggest a crucial role for *GZMK*-expressing populations in driving inflammatory responses in the CSF. Future studies should explore the role of the *GZMK*-expressing cells, including GZMK^+^CD8^+^ TEM and CD56^bright^ NK cells, in TBM pathogenesis.

Along with *GZMK*-expressing cells, microglia-like macrophages, CD4^+^ TM, and Treg populations were increased in the CSF of TBM patients, providing new insights into TBM immunopathogenesis. Microglia-like macrophages in the CSF of TBM adults highly expressed genes for Toll-like receptors (*TLR2*/*4*/*7*), which are involved in *Mtb* recognition and phagocytosis, as well as complement-activating components (*C1QA*/*B*/*C*) and genes related to interferon (*CXCL9*/*10*) and inflammatory (*APOE*, *APOC1*) responses. The CSF of TBM patients was predominantly comprised of specialized, tissue-resident and adaptive cell populations, while PBMCs showed relative enrichment of innate immune cell populations (e.g. monocytes) and naïve adaptive immune cells with high mobility. This distribution reflects the complementary and specific immunological roles of the two anatomical sites – blood cells circulate throughout the body and are prepared for rapid and systemic responses, whereas CSF cells are equipped to defend against pathogens on the frontline. However, the presence of all major cell clusters in both PBMCs and CSF, along with elevated inflammatory signaling in both compartments compared with a healthy donor, suggests that the immune response in blood partially reflects that at the site of infection.

In addition to increased inflammation, we observed reduced oxidative phosphorylation in multiple cell types in CSF compared with PBMCs. Similarly, in a *Mycobacterium bovis* (BCG)-induced TBM mouse model, scRNA-seq revealed decreased oxidative phosphorylation accompanied by increased activation of immune response and inflammatory pathways in diverse brain cell types (45). In TBM, immune cells undergo significant metabolic reprogramming, characterized by a shift from oxidative phosphorylation to glycolysis (46), potentially to meet the increased energy demands of inflammation. Alternatively, reduced oxidative phosphorylation could reflect mitochondrial dysfunction caused by elevated levels of reactive oxygen species generated during active immune responses (46). These metabolic and inflammatory changes in CSF highlight the complex interplay between energy metabolism and immune activation in driving TBM pathogenesis.

Although limited by sample number, our scRNA-seq study is consistent with findings from bulk RNA-seq (6), demonstrating elevated inflammatory responses and TNF signaling in PBMCs and CSF in TBM patients compared to a healthy control, and in a non-survivor compared to survivors. However, it also provides deeper insights by identifying the specific cell types and gene pathways associated with mortality. For instance, inflammatory signaling pathways were increased in multiple cell types but not in B cells, CD4^+^ naïve/TCM, or CD8^+^ naïve/TCM. Additionally, the apoptosis pathway was specifically elevated in highly inflammatory cells, such as GZMK^+^CD8^+^ TEM cells, monocytes, and microglia-like macrophages, and in highly cytotoxic GNLY^+^CD8^+^ TEM cells, in PBMCs and CSF of the non-survivor. While these results require further investigation, the identification of key cell types driving disease outcomes through scRNA-seq holds promise for significant advancements in biomarker discovery and targeted therapies for TBM.

In conclusion, our study demonstrates the potential of scRNA-seq for uncovering the cellular and molecular mechanisms underlying complex inflammatory responses in TBM. We successfully characterized the immune composition and gene expression profiles in PBMCs and CSF, revealing key differences between these compartments that may contribute to TBM mortality. Moreover, the scRNA-seq workflow and analysis pipeline that we established provides a valuable framework for future single-cell transcriptomic studies of blood and CSF to gain insights into the pathogenesis and identify potential treatment strategies for TBM and other brain infections.

## Acknowledgements

We would like to thank the doctors and nurses who recruited the patients into our study at the Hospital for Tropical Diseases, Ho Chi Minh City, Vietnam, and all participants. The study was funded by Wellcome Trust Intermediate Fellowship [206724/Z/17/Z to N.T.T.T.] and Wellcome Trust Investigator Award [110179/Z/15/Z to G.E.T., 222426/Z/21/Z to P.K.]. For the purpose of open access, the authors have applied a CC BY public copyright license to any Author Accepted Manuscript version arising from this submission. The funding body has no role in the design of the study, collection, analysis, interpretation of data, or in writing the manuscript.

## Authors’ Contributions

Conceptualization: T.T.B.T., L.C.G., P.K., N.T.T.T.; Supervision: P.K., N.T.T.T.; Methodology: T.T.B.T., L.C.G., L.N.H.T., L.T.H.N., D.D.A.T., H.D.T.N., L.H.V., V.T.N.H.; Funding acquisition: G.E.T., P.K., N.T.T.T.; Writing - Review & Editing: T.T.B.T., L.C.G., L.N.H.T., L.T.H.N., D.D.A.T., H.D.T.N., L.H.V., G.E.T., V.T.N.H., P.K., N.T.T.T.

## Abbreviations

TBM: Tuberculosis meningitis
scRNA-seq: Single-cell RNA-sequencing
CSF: Cerebrospinal fluid
PBMCs: Peripheral blood mononuclear cells
*Mtb*: *Mycobacterium tuberculosis*
CNS: Central nervous system

