## Supplement Figures for "Single-cell profiling of blood and cerebrospinal fluid in tuberculous meningitis"

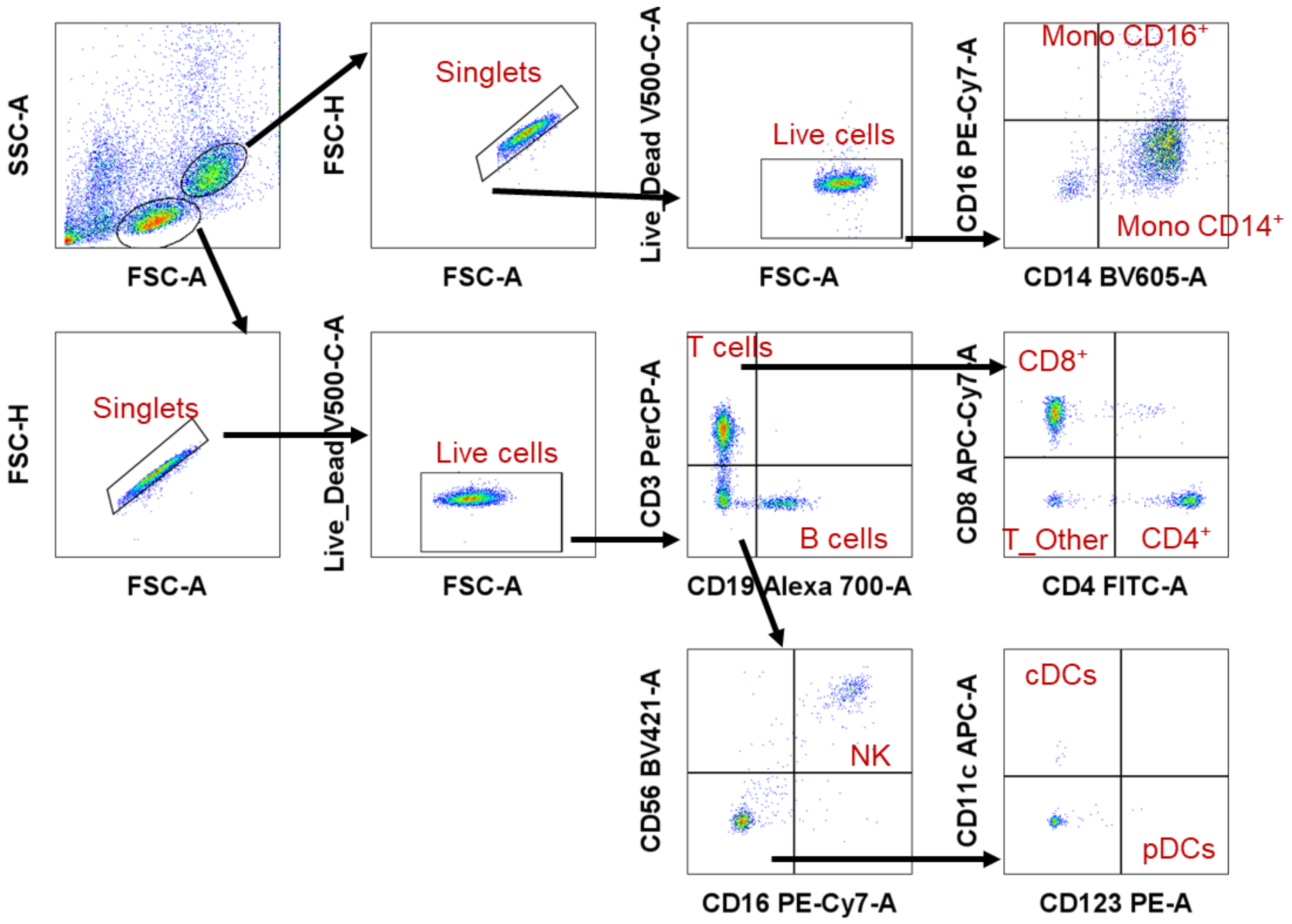


**Figure S1. Flow cytometry gating strategy**. Arrows depict the gating strategy. Mono: monocytes, $CD4^+$: $CD4^+$ T cells, $CD8^+$: $CD8^+$ T cells, T_Other: other $CD3^+$ T cells, DCs: dendritic cells cDCs: conventional dendritic cells, pDCs: plasmacytoid dendritic cells, NK: natural killer cells.

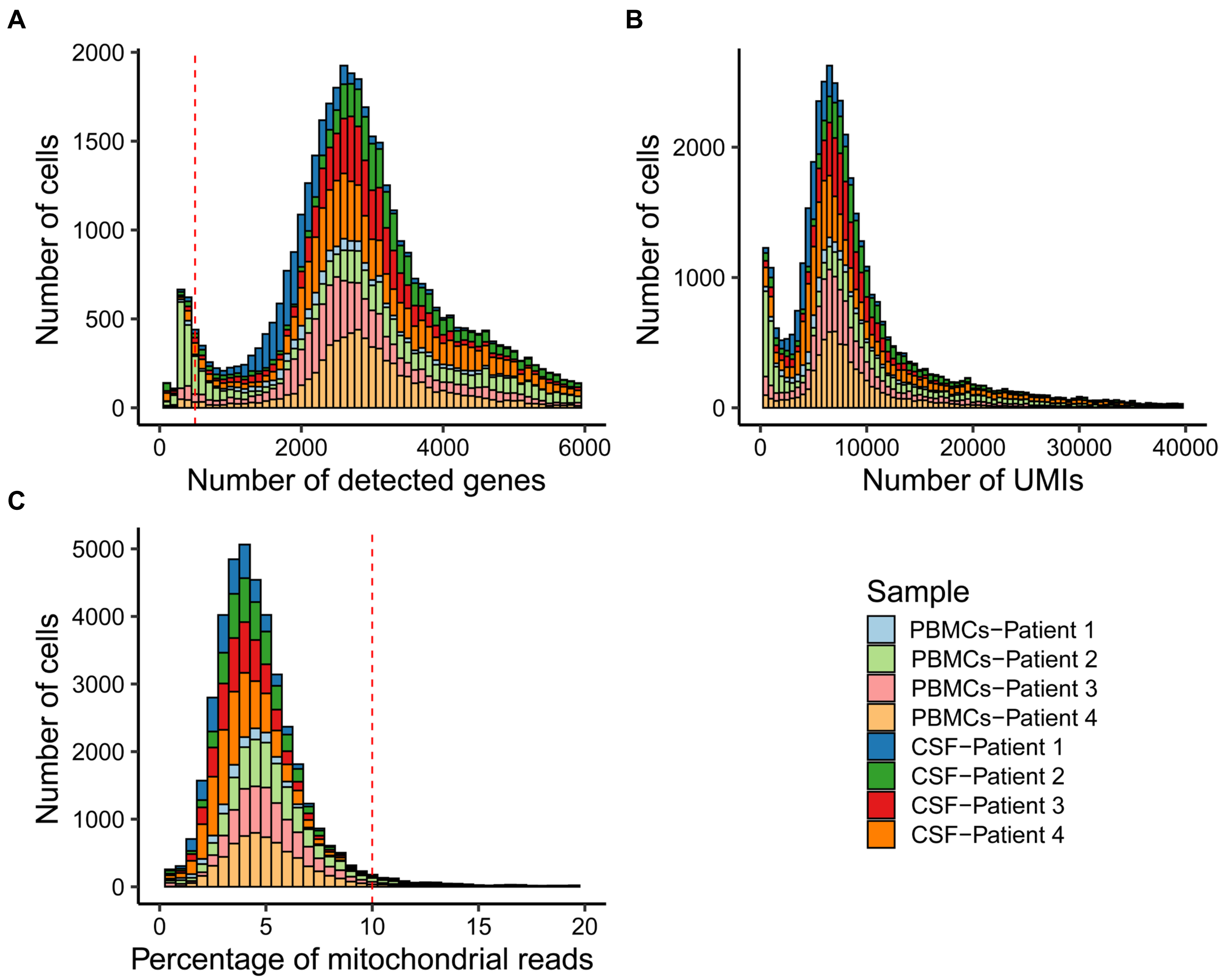


**Figure S2**. **Quality control of single-cell RNA-sequencing data**. Histograms of (**A**) the number of detected genes, (**B**) UMI counts, or (**C**) the percentage of mitochondrial reads for each single cell from PBMCs or CSF from TBM patients. Low-quality or dead cells, defined as those with more than 10% of reads mapping to mitochondrial genes and/or fewer than 500 detected genes, were discarded before further analysis, as indicated by the dashed red line.

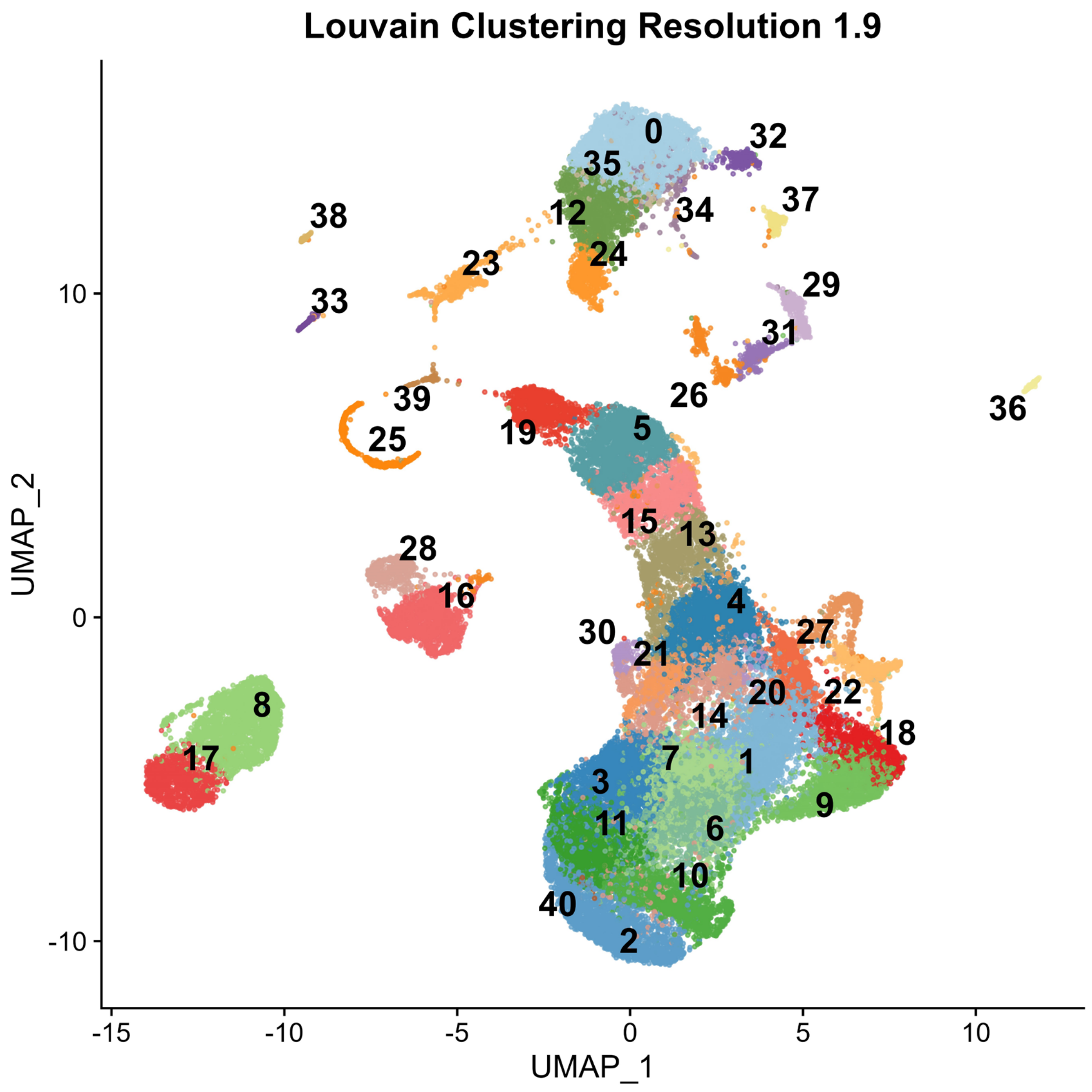


**Figure S3**. UMAP plot of Louvain clustering of combined PBMCs and CSF cells at a resolution of 1.9. Each cluster is color-coded and numbered.

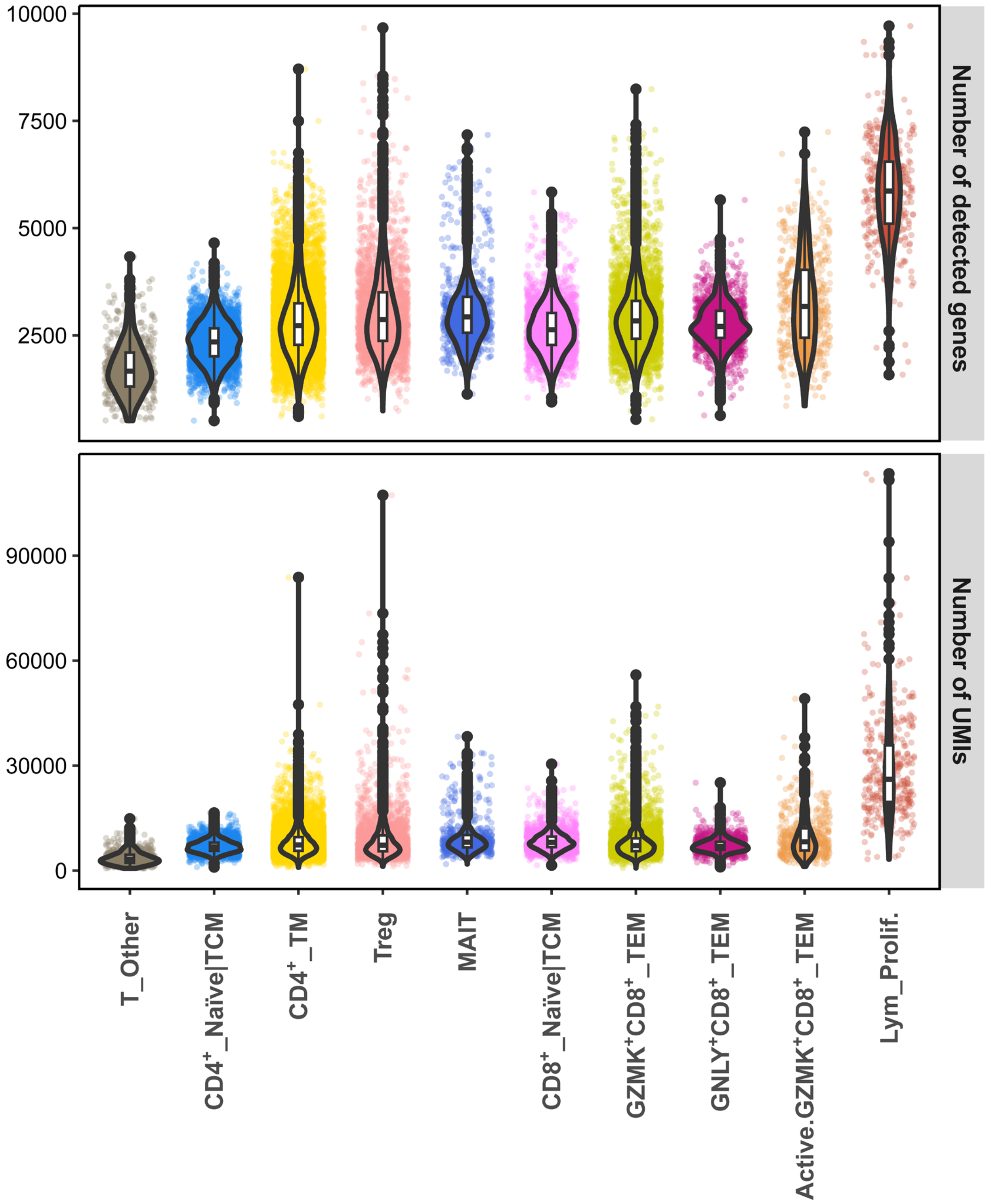


**Figure S4**. Low number of detected genes and total transcript (UMI [unique molecular identifier]) counts in T_Other compared to the rest of T cell types. Lym_Prolif.: proliferative lymphocytes, TCM: T central memory cells, TEM: T effector memory cells, TM: T memory cells.

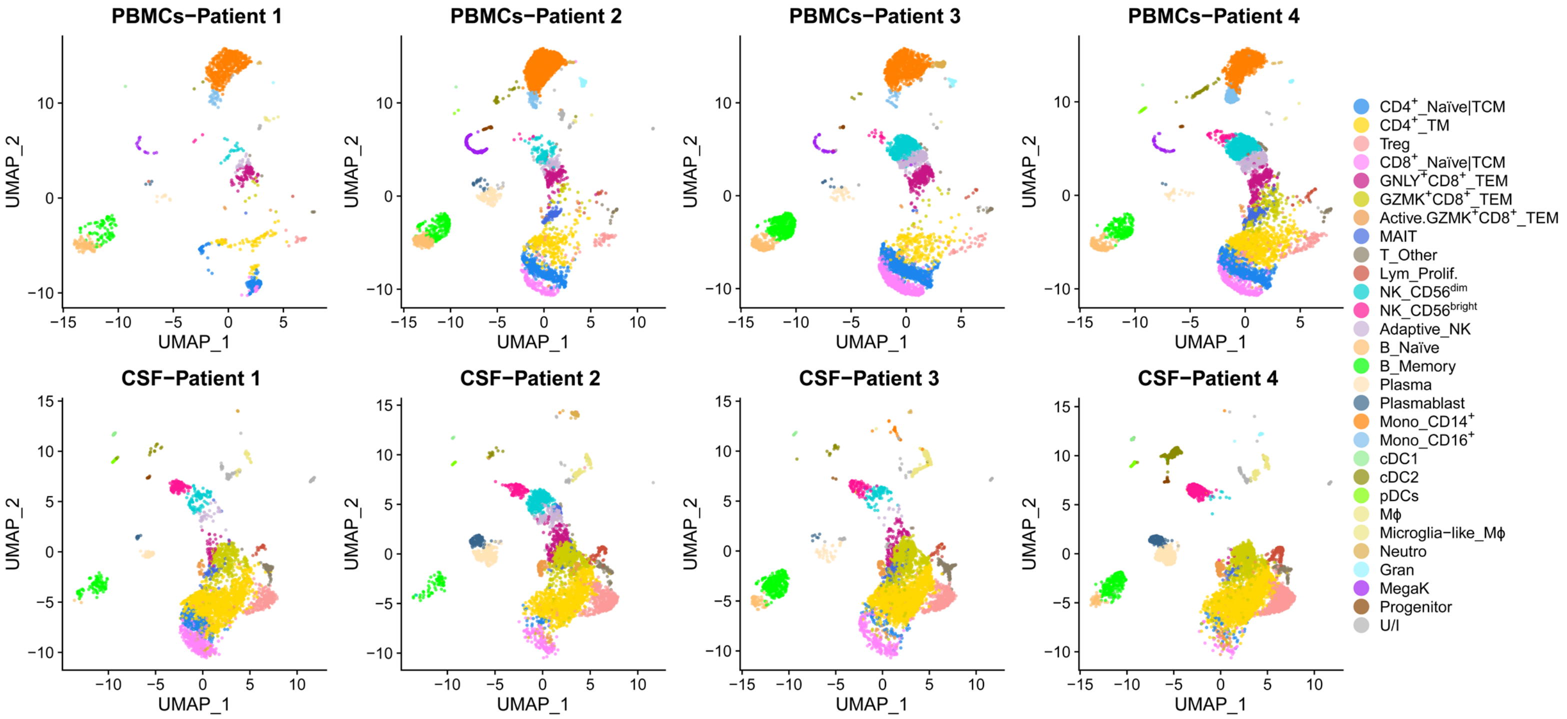


**Figure S5. Identification of cell subtypes (Level 2) in PBMCs and CSF stratified by donor.** Cell subtypes were annotated based on expert knowledge. cDC1: conventional DC1, cDC2: conventional DC2, Gran: mixed granulocytes, Lym_Prolif.: proliferative lymphocytes, MegaK: megakaryocytes, MΦ: macrophages, Neutro: neutrophils, pDCs: plasmacytoid DCs, TCM: T central memory cells, TEM: T effector memory cells, TM: T memory cells, U/I: unidentified.

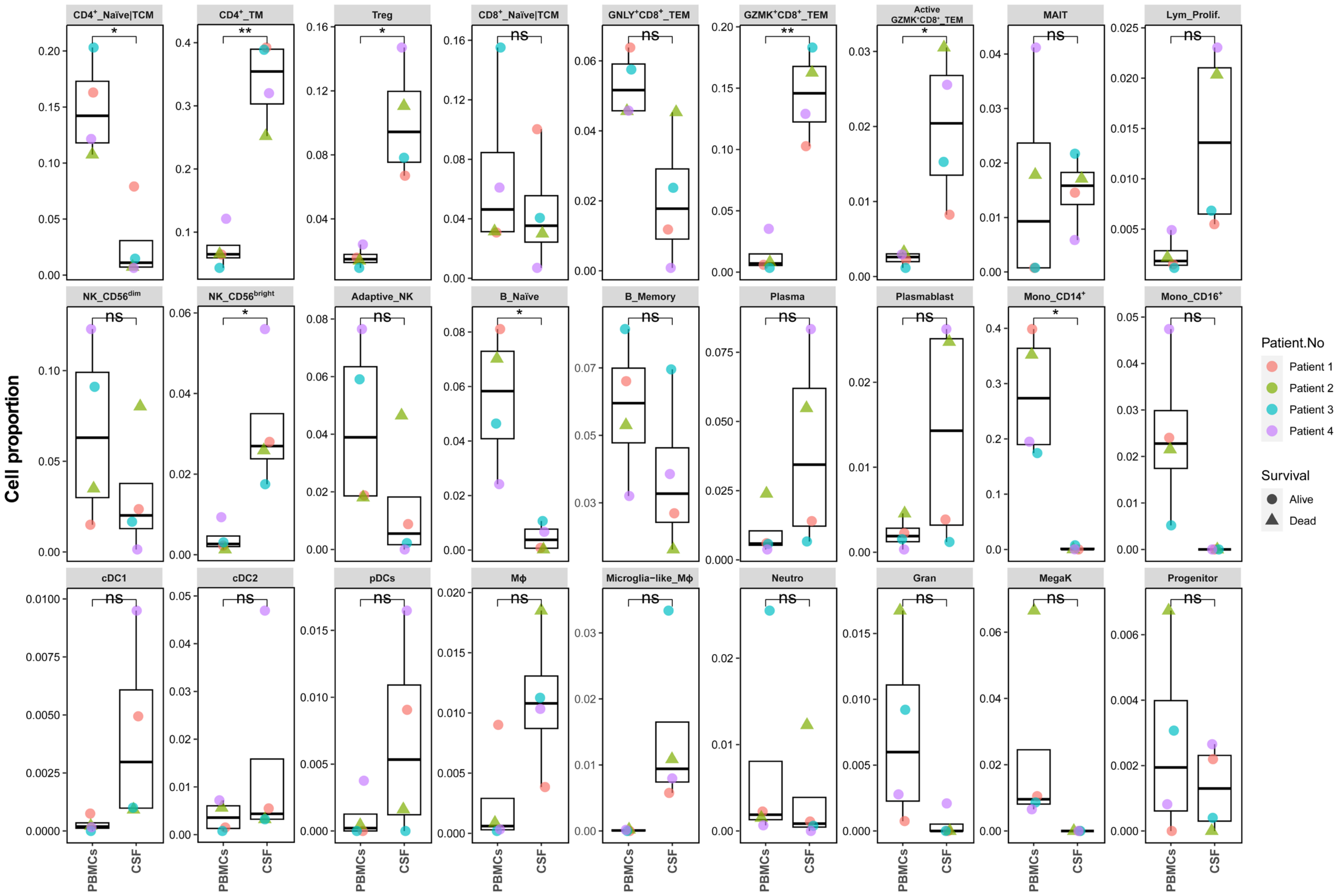


**Figure S6. Cell subtype proportions in PBMCs and CSF of individual patients.** Comparison of cell subtype proportions between PBMCs and CSF was performed using paired t-tests. *p < 0.05; **p < 0.01, ns: non-statistically significant difference. Adaptive_NK: adaptive-like NK, cDC1: conventional DC1, cDC2: conventional DC2, Gran: mixed granulocytes, Lym_Prolif: proliferative lymphocytes, MegaK: megakaryocytes, MΦ: macrophages, Neutro: neutrophils, pDCs: plasmacytoid DCs, TCM: T central memory cells, TEM: T effector memory cells, TM: T memory cells.

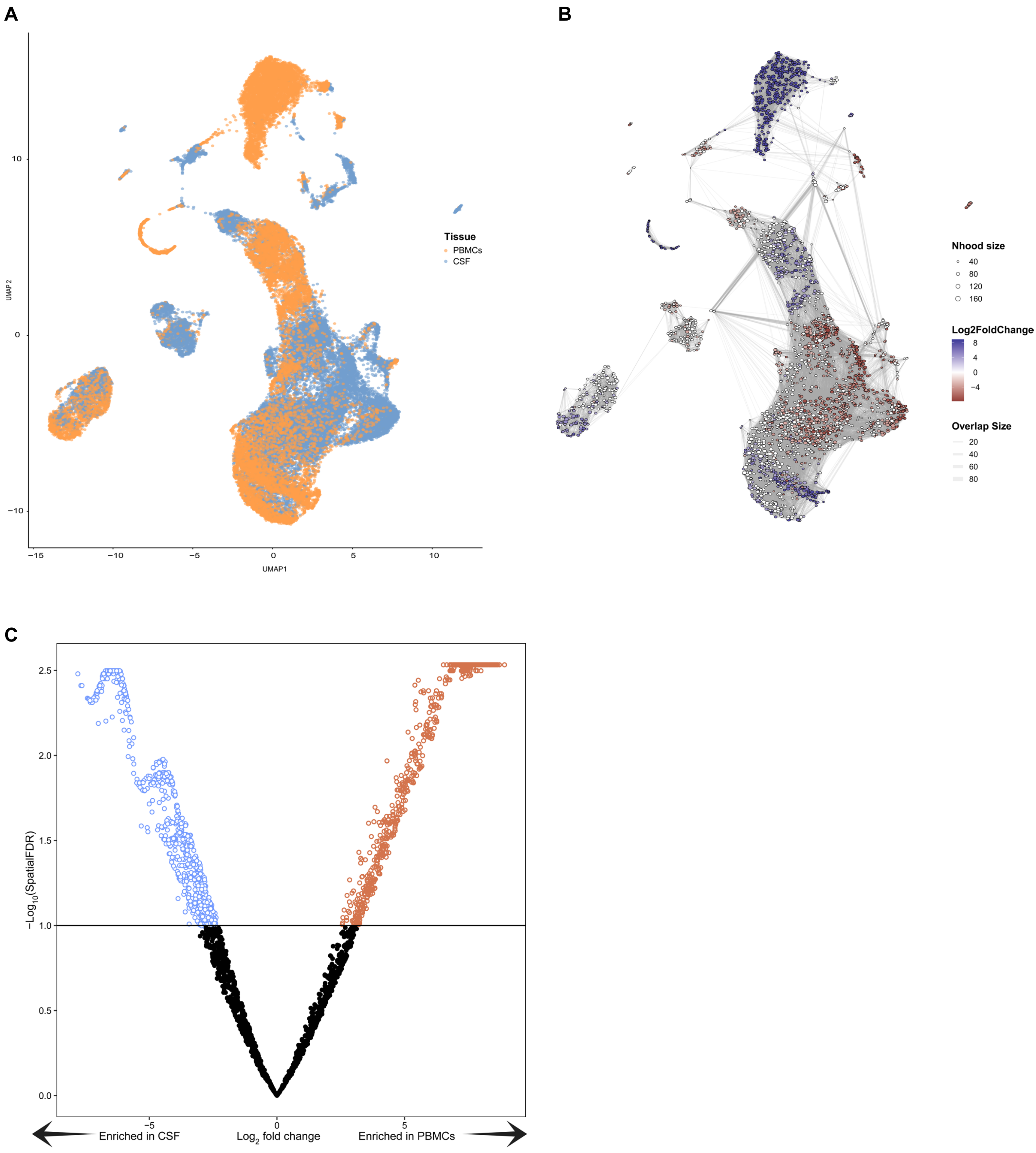


**Figure S7. k-NN weighted-network analysis to identify differentially abundant neighborhoods using miloR.** (**A**) UMAP plot of all cells colored by tissue of origin. (**B**) k-NN weighted-network analysis to identify differentially abundant neighborhoods using the miloR package. Nhood size indicates the number of cells in the neighborhood. (**C**) Volcano plot indicating differentially abundant (spatial FDR < 0.1) neighborhoods (n = 1845) between PBMCs and CSF.

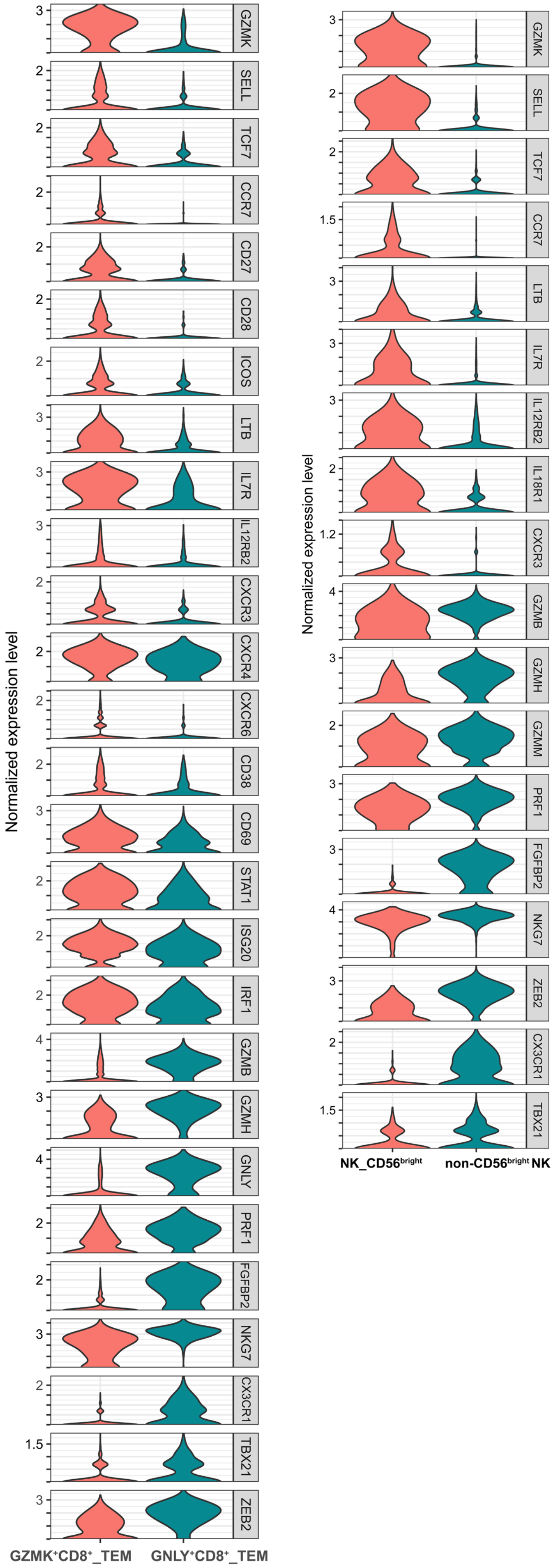


**Figure S8. Gene expression levels in CD8 and NK subpopulations.** Normalized expression levels of example differentially expressed genes between GZMK$^{+}$CD8$^{+}$ TEM and GNLY$^{+}$CD8$^{+}$ TEM, and between CD56$^{bright}$ NK and non-CD56$^{bright}$ NK were plotted.

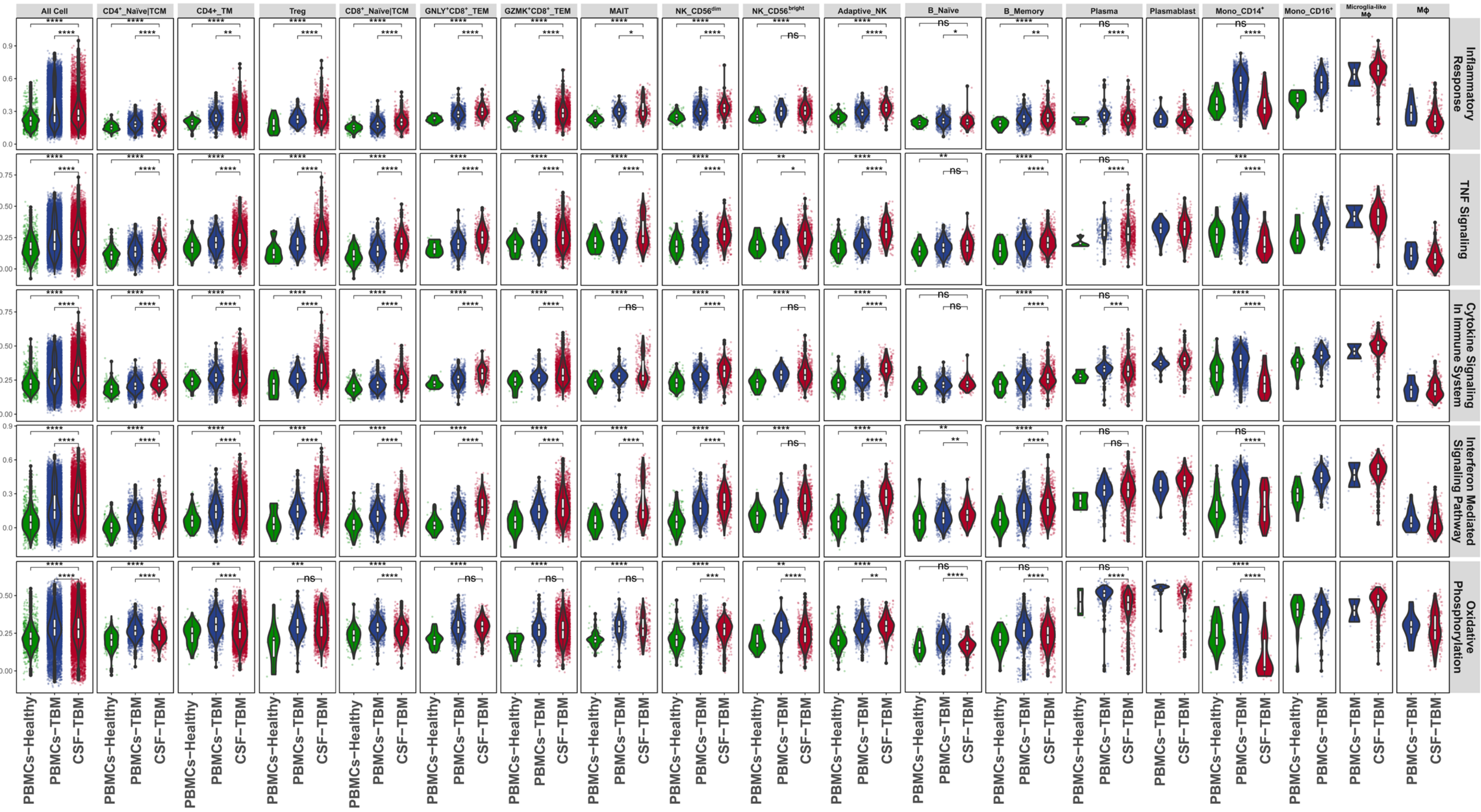


**Figure S9. Single-sample gene set enrichment analysis (ssGSEA).** Pathway normalized enrichment scores (NES) were calculated for individual cells using ssGSEA. Each dot represents one cell. Boxes indicate the median, and 25th and 75th percentiles, with whiskers extending to the minimum and maximum data points. Statistical comparison between groups was performed using the Wilcoxon rank-sum test. *p < 0.05, **p < 0.01, ***p < 0.001, ****p < 0.0001, ns: non-statistically significant difference.
